## Supplemental Material for "Cluster size determines internal structure of transcription factories in human cells"

(Dated: October 28, 2025)

### SIMULATION DETAILS

#### Multicolor bead polymer model

Chromatin filaments are coarse-grained into bead-and-spring polymers with $M$ monomers [1], each monomer having a diameter of $\sigma = 30$ nm, corresponding to a 3 kbp genomic region[2](a slightly different mapping from genomic to spatial distances would not affect the final results of our work [3]. There are three types of beads: $M_{nb}$ beads interacting repulsively with all TFs and so non-binding, $M_w$ beads weakly binding TFs, and $\{M_i\}$, $= 1, \ldots, n_c$ beads strongly binding to same-coloured TFs only; $n_c$ here denotes the number of TU colours. Thus the total number of chain beads is $M = M_{nb} + M_w + \sum_{i=1}^{n_c} M_i$. TFs are also modelled as spheres with the same diameter $\sigma$; they can either be in a binding (on) or non-binding (off) state. In the on state, TFs can bind strongly to TUs of the same color, while in the off state they do not bind. At every time interval, $t_s$, each TF can change its state becoming off from on, or vice versa, with rates $\alpha_{off}$ and $\alpha_{on}$, respectively. The total number of TFs is $N = \sum_{i=1}^{n_c} 2N_i$, where the number of TU and TF colours is the same and the factor of 2 takes into account the on/off counting (for each colour, $N_i$ TFs are initially set on, and $N_i$ initially off). Finally the total number of beads is $M + N$, the position of each monomer (either TFs or chain monomers) is denoted as $\mathbf{r}_i = (r_{ix}, r_{iy}, r_{iz}),\ i = 1, \ldots, M + N$, and the distance between the generic $i$-th and $j$-th beads is denoted as $r_{ij}$.

To ensure chain connectivity, two consecutive chain beads $i$ and $j = i + 1$ interact through the harmonic potential:

$$U_{harmonic} = -k_h \left(r_{ij} - \bar{r}\right)^2 \ ,$$

where $k_h = 100 k_B T/\sigma^2$ is the harmonic spring constant, $\bar{r}$ is the equilibrium spring distance set at $\bar{r} = 1.1\ \sigma$, $k_B$ is the Boltzmann constant, and $T$ is temperature. Chain stiffness is described via a bending (Kratky-Porod) potential depending on the position of every three consecutive chain beads as follows:

$$U_{KP} = \frac{k_B T l_P}{\sigma} \left[1 - \frac{\vec{s_i} \cdot \vec{s_j}}{|\vec{s_i}||\vec{s_j}|}\right] = k_{kp} \left[1 - \cos(\theta)\right] \ ,$$

where $i$ and $j = i + 1$ are neighbouring polymer beads, $\vec{s_i}$ is the tangent vector connecting the $i$-th and the $i+1$-th beads, $\theta$ is the angle formed by such tangents, and $l_P$ is the persistent length of the chain.

Any two beads $i$ and $j$ interact in general through a truncated and shifted Lennard-Jones potential:

$$U_{LJ} = \begin{cases} 4\epsilon_{ij} \left[\left(\frac{\sigma}{r_{ij}}\right)^{12} - \left(\frac{\sigma}{r_{ij}}\right)^{6} - c\right] & \text{if } r_{ij} < r_c \\ 0 & \text{otherwise} \end{cases} \ ,$$

with $c = \left(\frac{\sigma}{r_c}\right)^{12} - \left(\frac{\sigma}{r_c}\right)^{6}$, and $r_c$ is the interaction cut-off. Different cut-offs and interaction strengths are set depending on the chain bead type and colour, and TF colour, of the two interacting beads. We use $r_c = 1.8\sigma$ and $\epsilon_{ij} = 8k_BT$ for strong binding between TUs and on TFs of the same colour, $r_c = 1.8\sigma$ and $\epsilon_{ij} = 3k_BT$ for weak binding between chain beads and all on TFs, and $r_c = 2^{1/6}\sigma$ (purely repulsive potential) and $\epsilon_{ij} = k_BT$ for non-binding interactions between non-binding polymer beads and TFs, between pairs of TFs, between pairs of TUs, and between pairs of different-colour TU and TF beads.

A TU bead is considered transcriptionally active when it is bound to at least one same-colored TF, i.e. when their distance is smaller than a fixed threshold. In order to be quite conservative about the evaluation of the transcriptional activity, here we consider $2.25\sigma$ as such a threshold. However, we checked that varying it within reasonable limits (we tested $1.8\sigma$ and $1.1\sigma$, see Fig. S11) does not affect the output transcriptional profiles in any significant way.

The time evolution of the entire systems is described by the following system of $3(M + N)$ Langevin equations

$$m_l \frac{d^2 r_{l\beta}}{dt^2} = -\nabla U_l - \gamma_l \frac{dr_{l\beta}}{dt} + \sqrt{2k_BT\gamma_l}\ \eta_{l\beta}(t) \ , \tag{1}$$

where $l$ is the bead index running from 1 to $M + N$, $\beta = x, y, z$ is the dimension index, $m_l$ and $\gamma_l$ respectively are the mass and friction coefficient associated to the $l$-th bead, $U_l$ is the total potential experienced by the $l$-th bead, and

$\eta_{l\beta}$ are a set of zero-mean delta-correlated stochastic white noises, in symbols

$$\langle \eta_{l\beta}(t) \rangle = 0 \ , \qquad \langle \eta_{l\beta}(t)\eta_{l'\beta'}(t') \rangle) = \delta_{ll'}\delta_{\beta\beta'}\delta(t-t') \ ,$$

where $\delta_{ll'}$ and $\delta_{\beta\beta'}$ are two Kronecker deltas and $\delta(t-t')$ is a Dirac delta. For the sake of simplicity, we set the mass and friction coefficient equal for all beads, $m_l \equiv m$ and $\gamma_l \equiv \gamma$ respectively.

#### Simulation units and system constants

As typically done in the literature [4], we introduce the Lennard-Jones time $\tau_{LJ} = \sigma\sqrt{m/\epsilon}$, with $\epsilon = k_B T$ the energy unit, and the Brownian time $\tau_B = \sigma^2/D$, with $D = k_B T/\gamma$ the diffusion coefficient of a single bead, representing the characteristic time a bead takes to diffuse across its own diameter. Our simulations are performed by setting mass $m$, diameter $\sigma$, temperature $T$, and Boltzmann constant $k_B$ to one. We also set the friction coefficient $\gamma = 1$, so that $\tau_{LJ} = \tau_B$, and $k_{kp}/k_B T = l_P = 3\sigma$, making the fibre relatively flexible. The simulation reduced units are then $\sigma$ for the length, $\epsilon$ for the energy, and $\tau_B$ for the time.

Protein switching is implemented by stochastically changing the TF state of all colours every $t_s = 10^2\tau_B$ with probability such that the switching off rate is $\alpha_{off} = 10^{-5}\ \tau_B^{-1}$. Both in the toy model and chromosome simulations the switching on rate is instead $\alpha_{on} = \alpha_{off}/4$, so that in the steady state for each colour the average number of off TFs is 4 times the number of the on ones.

The system is free to evolve in a cubic simulation box with periodic boundary conditions of side $L = 173\sigma$ for the toy model, $L = 570\sigma$ for HUVEC and GM12878 HSA14, $L = 500\sigma$ for HUVEC HSA18 and $L = 556\sigma$ for HUVEC HSA19 so that in all cases the systems is in the dilute regime and the TU density is the same in all cases. In all toy model cases the initial number of TFs per color is set at 25, except for the edited one where the initial number of yellow on and off TFs is reduced by 30%, from 25 to 17.

#### Mapping to physical units

Conversion from simulation to physical units is readily obtained. Using the realistic values of bead diameter $\sigma = 3\ 10^{-8}\ m$ (representing also chain thickness), room temperature $T = 300\ K$ (giving $\epsilon \simeq 4.14\ 10^{-21}\ J$) and Stokes friction coefficient for spherical beads $\gamma = 3\pi\eta_{sol}\sigma$, with nucleoplasm viscosity $\eta_{sol} = 10 - 100\ cP$, one finds that $\tau_{LJ} = \tau_B = \sigma^2/D = 3\pi\eta_{sol}\sigma^3/\epsilon \simeq 0.6 - 6\ 10^{-3}\ s$.

#### Simulation details

We use the LAMMPS (Large-scale Atomic/Molecular Massively Parallel Simulator) software package [5] to simulate the system. For computational efficiency, hydrodynamic interactions are neglected. The system of equations Equation 1 is integrated using the velocity Verlet algorithm. The timestep is set to $\Delta t = 0.01\tau_B$.

To begin simulations, the chromatin fibre is initialised as a random walk while TFs are placed randomly within the simulation box. Overlapping beads in the initial conformation are relaxed evolving the system for a few timesteps with a soft potential between beads. The system is then thermalised for $10^5\ \tau_B \simeq 60 - 600\ s$ for the toy model and for $4\ 10^4\ \tau_B \simeq 24 - 240\ s$ for whole chromosomes, considering only repulsive interaction between all couples of beads. Finally, the system is evolved including all attractive interactions for $8\ 10^5\ \tau_B \simeq 480 - 4800\ s$ for the toy model and for $2\ 10^5\ \tau_B \simeq 120 - 1200\ s$ for chromosomes. For each considered case, we run at least 100 independent simulations.

#### Toy model TU color sequences

We report here the TU colour sequences (bead indexes) of all cases considered for the toy model. All other beads are weakly-binding, apart from the non-binding case analysed in Figure S4 and Figure S7 in which all other beads are non-binding, and which has the same TU color sequence as the reference case.

- Reference case.
  Red TU beads: 60, 210, 300, 360, 390, 480, 630, 660, 720, 990, 1080, 1110, 1170, 1200, 1350, 1380, 1410, 1440, 1530, 1560, 1590, 1620, 1650, 1800, 1890, 2190, 2310, 2460, 2520, 2670, 2730, 2760, 2790, 2910;

Yellow TU beads: 30, 120, 570, 930, 1050, 1140, 1230, 1260, 1290, 1320, 1710, 1770, 1860, 1920, 1950, 1980, 2010, 2040, 2070, 2130, 2160, 2220, 2340, 2370, 2400, 2430, 2550, 2580, 2610, 2700, 2970;
Green TU beads: 90, 150, 180, 240, 270, 330, 420, 450, 510, 540, 600, 690, 750, 780, 810, 840, 870, 900, 960, 1020, 1470, 1500, 1680, 1740, 1830, 2100, 2250, 2280, 2490, 2640, 2820, 2850, 2880, 2940, 3000;

- $n$-mutant cases. Same TU color sequence as reference case except for yellow→red mutations of the following TU beads:
  2010 for 1-mutant case;
  1980 and 2010 for 2-mutants case;
  1950, 1980 and 2010 for 3-mutants case:
  1950, 1980, 2010 and 2040 for 4-mutants case;

- 6-pattern case.
  Red TU beads: 30, 60, 90, 120, 150, 180, 570, 600, 630, 660, 690 ,720, 1110, 1140, 1170, 1200, 1230, 1260, 1650, 1680, 1710, 1740, 1770, 1800, 2190, 2220, 2250, 2280, 2310, 2340, 2730, 2760, 2790, 2820, 2850, 2880;
  Yellow TU beads: 210, 240, 270, 300, 330, 360, 750, 780, 810, 840, 870, 90, 1290, 1320, 1350, 1380, 1410, 1440, 1830, 1860, 1890, 1920, 1950, 1980, 2370, 2400, 2430, 2460, 2490, 2520, 2910, 2940, 2970, 3000;
  Green TU beads: 390, 420, 450, 480, 510, 540, 930, 960, 990, 1020, 1050, 1080, 1470, 1500, 1530, 1560, 1590, 1620, 2010, 2040, 2070, 2100, 2130, 2160, 2550, 2580, 2610, 2640, 2670, 2700.

- 1-pattern case.
  Red TU beads: 30, 120, 210, 300, 390, 480, 570, 660, 750, 840, 930, 1020, 1110, 1200, 1290, 1380, 1470, 1560, 1650, 1740, 1830, 1920, 2010, 2100, 2190, 2280, 2370, 2460, 2550, 2640, 2730, 2820, 2910, 3000;
  Yellow TU beads: 60, 150, 240, 330, 420, 510, 600, 690, 780, 870, 960, 1050, 1140, 1230, 1320, 1410, 1500, 1590, 1680, 1770, 1860, 1950, 2040, 2130, 2220, 2310, 2400, 2490, 2580, 2670, 2760, 2850, 2940;
  Green TU beads: 90, 180, 270, 360, 450, 540, 630, 720, 810, 900, 990, 1080, 1170, 1260, 1350, 1440, 1530, 1620, 1710, 1800, 1890, 1980, 2070, 2160, 2250, 2340, 2430, 2520, 2610, 2700, 2790, 2880, 2970;

- single-colour case. TU beads: 30, 60, 90, 120, 150, 180, 210, 240, 270, 300, 330, 360, 390, 420, 450, 480, 510, 540, 570, 600, 630, 660, 690, 720, 750, 780, 810, 840, 870, 900, 930, 960, 990, 1020, 1050, 1080, 1110, 1140, 1170, 1200, 1230, 1260, 1290, 1320, 1350, 1380, 1410, 1440, 1470, 1500, 1530, 1560, 1590, 1620, 1650, 1680, 1710, 1740, 1770, 1800, 1830, 1860, 1890, 1920, 1950, 1980, 2010, 2040, 2070, 2100, 2130, 2160, 2190, 2220, 2250, 2280, 2310, 2340, 2370, 2400, 2430, 2460, 2490, 2520, 2550, 2580, 2610, 2640, 2670, 2700, 2730, 2760, 2790, 2820, 2850, 2880, 2910, 2940, 2970, 3000;

### Human chromosome sequences

We detail here the procedure used to build the multicolour polymer bead sequences of the chromosomes. The procedure consists of two steps:

1. *Extraction of the bead sequence from experimental data for a single cell line.* ChIP-seq for H3K27ac data and DNase-hypersensitivity (DHS) data for the cell line of interest are used to determine the weakly-binding and TU bead positions along the chain, respectively.

   Concerning H3K27ac, two different unfiltered and control data sets are first downloaded from ENCODE [6] as .bam files and then combined into a main and control .bed files using bedtools [7]. The tool epic [8] is then employed to identify experimental peaks and produce the final .bed H3K27ac peak file. Peak identification by the epic tool works as follows: first the tool resolves main and control data at fixed resolution (in our case 200 bp); then it compares each resolved region of main and control data and assigns a peak if the amount of ChIP signal in the main data is significantly higher than the amount in the control data using as discriminant criterion the false discovery rate cutoff (in our case 0.05).

   Concerning DHSs, data are first downloaded from ENCODE as .bam files and, as before, two different data sets are taken into account. The peaks from alignment results are then identified and combined through the MACS [9] tool as for the H3K27ac data and cast into the human chromosome size through the bedGraphToBigWig conversion tool [10], the latter action being performed using as a reference for chromosome size the common hg19.chrom.sizes file [11]. The final DHS peak .bed file is then produced by considering only significant peaks (in our case $-\log_{10}(peak\ value) \geq 30$).

Finally the bead polymer sequence is produced by casting the .bed H3K27ac and DNase-hypersensitivity files into the cell line sequence as prescribed by the chosen polymer bp resolution.

2. *Specific cell line multicolour sequence generation.* The actual multicolour bead polymer sequence for a specific cell line is generated by combining bead sequences for the cell of interest and for the stem cell H1-hESC generated as prescribed by the previous step. Specifically, weakly-binding and TU bead attribution is performed by looking first at the specific cell bead sequences: if a bead contains an H3K27ac peak and no DHS peaks, it is marked as weakly-binding, while DHS peaks are marked as TUs. TU colours are instead determined by combining the specific cell line and H1-hESC TU bead sequences: if a TU appears in both the specific and H1-hESC cell, it is marked as housekeeping, while if a TU appears in the specific cell line sequence only it is marked as specific.

### HiP-HoP simulations of HSA 14 in HUVEC

In the main text, we present results regarding the mixing-demixing transition using the *highly predictive heteromorphic polymer model* or HiP-HoP [12] model (main figure 8). We present here the details, parameters, and input data of this model. In the HiP-HoP model the chromatin fibre is heteromorphic, meaning that its thickness is not constant. Indeed, in order to catch the less accessible chromatin structure of regions poor in H3K27ac marks [12], the polymer segments without this mark are modelled as crumpled by adding some springs. In particular, if a triplet of beads $i$, $i+1$ and $i+2$ do not include H3K27ac marks, a spring is applied between next-nearest neighbour beads ($i$ and $i+2$) and it is described by the following harmonic potential

$$V_{\mathrm{HARM}} = K_{\mathrm{HARM}}(r - R_{\mathrm{HARM}})^2,$$

with $K_{\mathrm{HARM}} = 200\epsilon/\sigma^2$ being the spring constant and $R_{\mathrm{HARM}} = 1.1\sigma$ the equilibrium bond length. In the HiP-HoP model we include a loop extrusion mechanism which is considered to be performed by the cohesin complex[13–15]. The action of cohesin rings is modelled by inserting additional spring bonds between beads; the pair of beads bonded by these springs is then moved outwards, i.e. from $(i, i+3) \to (i-1, i+4) \to (i-2, i+5)$ etc., to extrude a loop. Extruders (i.e. cohesin bonds) are loaded on the polymer at random $i, i+3$ positions with rate $k_{\mathrm{on}}$ and they unbind with rate $k_{\mathrm{off}}$. Their positions are moved at rate $k_{\mathrm{ex}}$. The spring potential is given by

$$U_{\mathrm{EXTR}}(r_{i,j}) = U_{\mathrm{WCA}}(r_{i,j}) + K_{\mathrm{EXTR}}(r_{i,j} - r_0)^2,$$

where $K_{\mathrm{EXTR}} = 40k_BT/\sigma^2$ is the bond strength and $r_0 = 1.5\sigma$ is the equilibrium spring length. The two ends of each extruders move independently and each of them is halted either either when it collides with the end of another extruder, or when it reaches a CTCF site whose direction is opposite to its direction of travel. If an extruder is halted on one end, the position at the other end keeps moving, extruding the loop. The loop extrusion dynamics is obtained by using python script using the LAMMPS library.

Differently from the previously discussed polymer model, four species of transcription factors (TFs), represented by spheres of the same size as the polymer beads, are inserted in the model, two active, as in the previous model, and two new inactive species. The two active TF types correspond again to general active complexes in charge of transcription of genes [16] specific of HUVEC cells and of genes shared by HUVEC and H1-hesc cells respectively. The two new inactive TF species model polycomb repressive complexes (PRCs) [17] and heterochromatin proteins (such as HP1)[18]. The interactions between TFs and polymer beads are described by the same shifted and truncated Lennard-Jones potential discussed before. The new Inactive TFs experience the following potentials: polycomb-like proteins are highly attracted ($\varepsilon = 8\,k_BT$) to chromatin beads enriched in H3K27me3, while heterochromatin TFs experience a strong attraction ($\varepsilon = 8\,k_BT$) to chromatin beads containing H3K9me3 marks.

Finally, to facilitate the phase separation between euchromatin and heterochromatin, an additional weak chromatin-chromatin attraction is inserted between beads not provided with H3K27ac marks. This last potential is described by the shifted and truncated Lennard-Jones potential $V_{LJ/cut}$ with $\varepsilon = 0.4\,k_BT$. As in previous model, during the simulations TFs switch back and forth between a binding and a non-binding state with at rate $k_{\mathrm{switch}}$[19]. This represents post-translation modifications which alter the protein-DNA binding affinity (e.g. phosphorylation). When in the non-binding state the interaction between proteins and chromatin binding sites revert to the WCA potential.

We simulate chromosome 14 in HUVEC cells, which, by using a resolution of $3\,\sigma$, is composed of 35,784 beads. The polymer is immersed in a cubic simulation box of side $84.5\,\sigma$ with periodic boundary conditions. This results in a chromatin density of $\sim 60\, bp/\sigma^3$, which is the same order of magnitude as *in vivo*, once mapped to real units (see "Mapping simulation units to real units"). The number of active TFs is set equal to 1200 (half in the binding state and half in the non-binding state), both for HUVEC-specific TFs and for TFs interacting with DHS sites found both in HUVEC and H1-hESC cells. We insert then 400 polycomb-like TFs (half in the binding state) and 1000 HP1-like proteins (again, half binding the chromatin fibre).

As initial condition, we generate the polymer as a random walk within the simulation box and we place TFs in random positions. We let the system to relax, initially inserting soft potentials to displace overlapping beads and then by using WCA steric interactions for non-bonded beads. A further equilibration is performed after inserting the additional crumpling springs between proper beads. At this point we create the initial configuration for HiP-HoP simulations by assigning a type to each polymer bead. The bead type depends on the combination of DHS peaks, H3K27ac, H3K27me3 and H3K9me3 marks present in the genome region covered by the bead. HiP-HoP simulations are performed by inserting 1000 extruders, which provide an extruder density of $\sim 10$ extrudes $Mbp$, comparable to the one observed *in vivo*.

We run 40 independent HiP-HoP simulations, each for a total time $5 \times 10^5\, \tau_B$. We discard the first $10^5 \tau_B$ (by which the system reaches a steady state), so that the analysis is performed on the last $4 \times 10^5 \tau_B$.

As in our previous model, in HiP-HoP simulations lengths are given in multiples of the polymer bead diameter $\sigma$. As we assume each bead contains $\sim 3\, kbp$, we set $\sigma = 30\, nm$. The simulation time unit is defined as the Lennard-Jones time $\tau_{LJ}$. To map the latter, we consider the other two characteristic times of the system: the Brownian time $\tau_B$ and the decorrelation time $\tau_{dec}$.

As discussed above and in the main text, TFs switch back and forth between a binding and non-binding state. The other simulation rates are related to loop extrusion and are set as: $k_{on} = 2 \times 10^{-2}\, \tau_B^{-1}$, $k_{off} = 5 \times 10^{-5}\, \tau_B^{-1}$, and $v_{ex} = \sigma \cdot k_{ex} = 8 \times 10^{-3}\, \tau_B^{-1}$ where $v_{ex}$ is the extrusion velocity. While these parameters were chosen to optimise simulation performances, the extruder density and the processivity ($\lambda = v_{ex}/k_{off}$ which in our case is $\lambda = 160kbp$) are compatible with the ones found in literature [13]. The simulation type of each chromatin bead depends on the presence of epigenetic marks and DNA accessibility data for the genomic region covered by the bead. Three epigentics marks are used: H3K27ac, H3K27me3 and H3K9me3 which are associated to transcriptionally active euchromatin, facultative heterochromatin and constitutive heterochromatin respectively. Their ChIP-seq profiles for the HUVEC cell line are obtained from the ENCODE database. The presence of H3K27ac marks is used to place the additional crumpling springs, while H3K27me3 and H3K9me3 peaks determine the chromatin binding sites for polycomb-like and heterochromatin proteins respectively. DNA accessibility is inferred from DHS data: DHS peaks in common between HUVEC and H1-hESC cells are used to know the position of active transcription factors which are cell generic, while DHS peaks present only in HUVEC cells determine the positions of HUVEC-specific active transcription factors. To include loop extrusion, CTCF binding sites have to be inserted in order to know where extruders halt. These are obtained from ChIP-seq peaks for CTCF which overlap with Rad21 ChIP-seq peaks. We then found the directionality of each site by locating its underlying binding motif. The center of the peak is mapped to a specific polymer bead that is so identified as a CTCF bead. To take into account cell-to-cell variability, in each repeated simulation we include only a part of CTCF beads, choosing them stochastically with a probability based on the peak height. All the data used as input are taken from the ENCODE database and are publicly available.

### SUPPLEMENTARY FIGURES

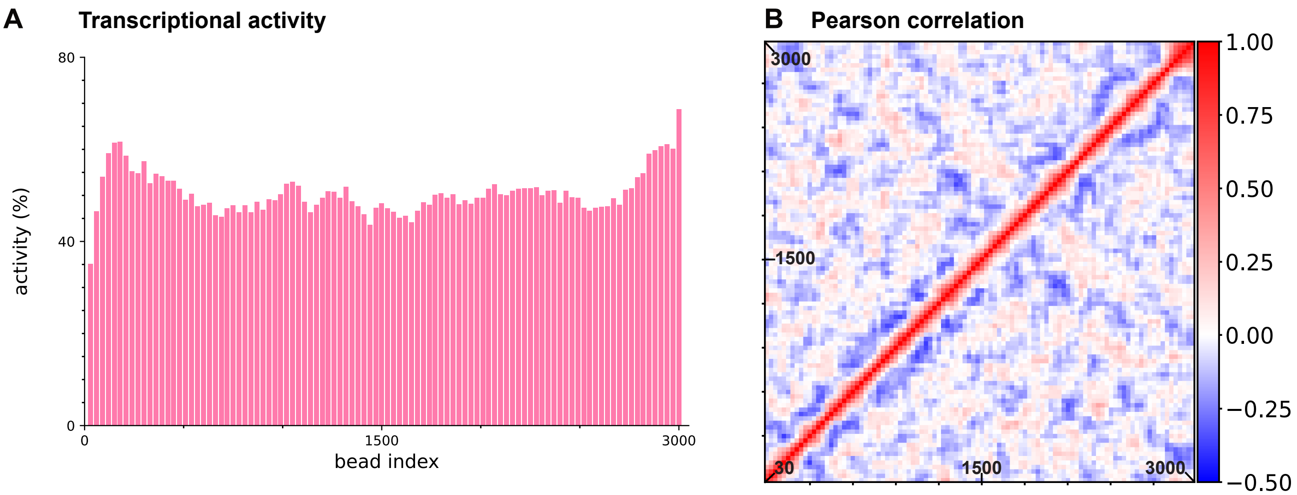


FIG. S1. **Random string with just one colour.** TUs in the random string considered in the main text are coloured randomly yellow, red, or green; here, instead, every TU has the same colour. Averages are obtained from 100 runs, each lasting $8\ 10^5\tau_B$. **(A)** The transcriptional activity profile is much flatter than that obtained with the 3-colour version. **(B)** The Pearson correlation matrix also lacks red blocks along the diagonal.

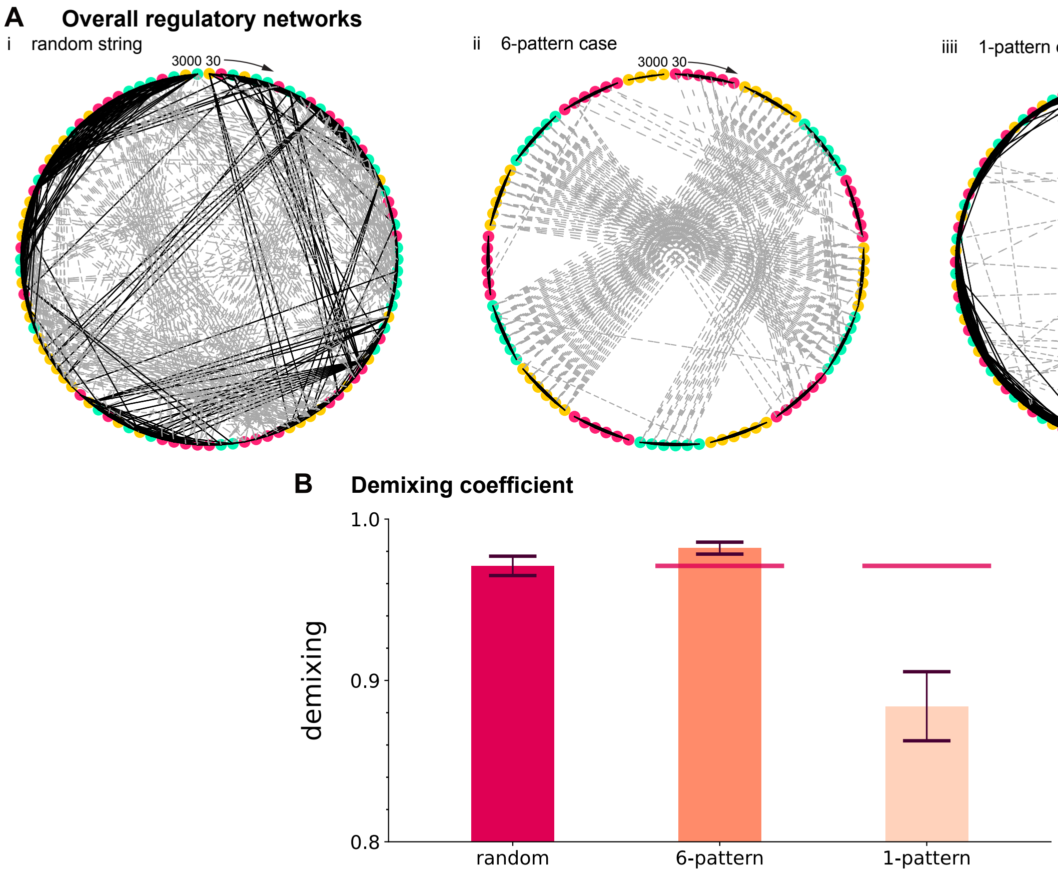


FIG. S2. **Complete regulatory networks, and demixing coefficients for toy strings. (A)** Networks involving positive (black) and negative (dashed grey) correlations between all coloured beads are presented, for the models considered in the main text in Fig. 5. **(i)** The random string. The network is complex and entails both positive and negative correlations. **(ii)** The 6-pattern repeat. There are many black edges within repeats and virtually none between repeats, many grey edges between repeats, and few edges between TUs with different colours. **(iii)** The 1-pattern repeat. There are few long black edges. **(B)** Average of $\theta_{\rm dem}$ (error bars: weighted SD) for various toy strings. Red lines: values for random string. Compared to the random case, 6-pattern is more demixed, and the 1-pattern less demixed. As discussed in the main text, same-colour and different-colour correlations diverge from each other in the 6-pattern case, and roughly overlap in the 1-pattern.

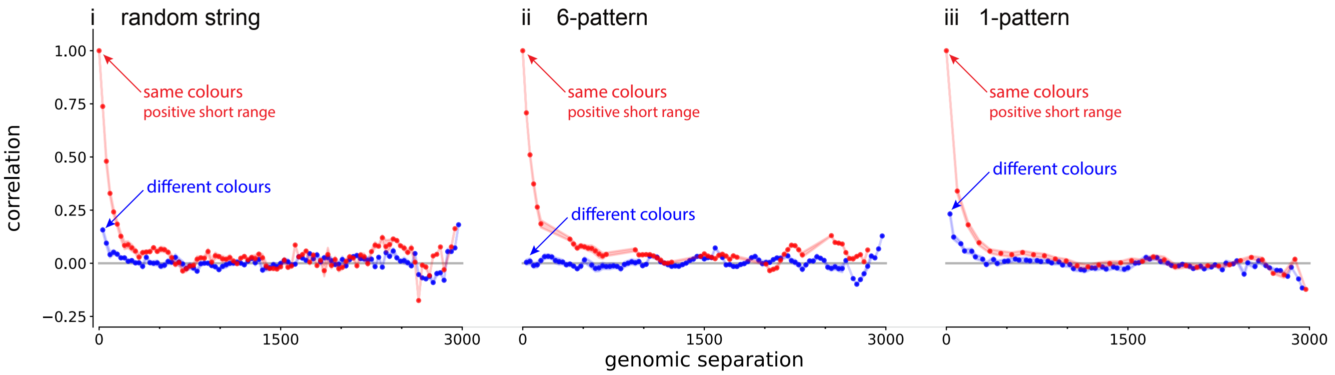


FIG. S3. **Activity correlations found in toy chromosomes without weakly-binding beads.** Correlation values (shading shows +/-SD, and is usually less than line/spot thickness) at fixed genomic distance are taken from super-/sub-diagonals of Pearson matrices. Red dots give mean correlations between TUs of same colour, and blue dots those between TUs of different colours. **(i)** Random string. **(ii)** 6-pattern string. **(iii)** 1-pattern string with non-binding beads instead of weakly-binding ones.

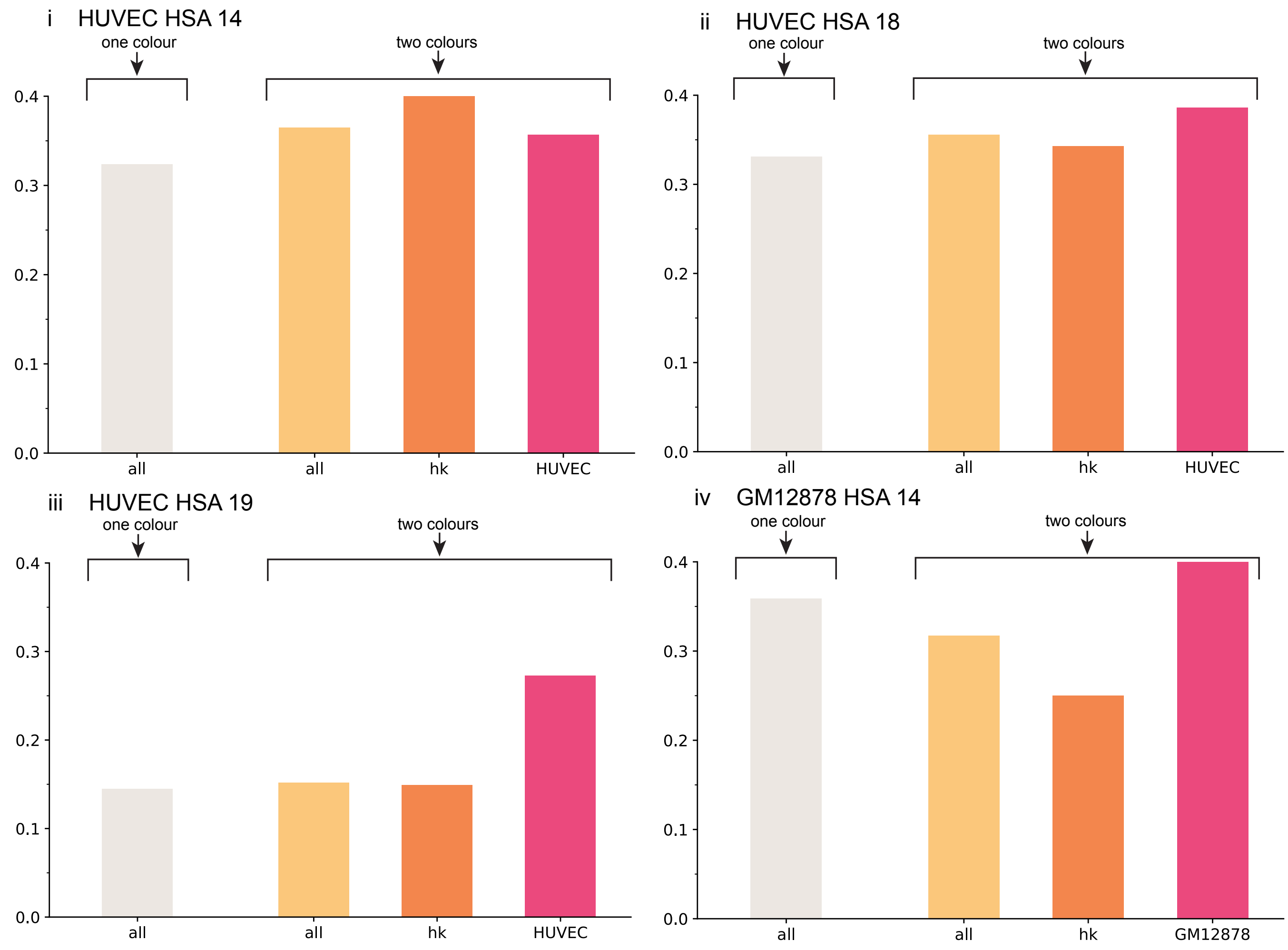


FIG. S4. **Spearman's rank-correlation coefficients determined by comparing activity data obtained from simulations (one- and two-colour) and GRO-seq for chromosomes and cell types indicated.** Grey bars: one-colour cases. Coloured bars: two-colour cases where values are for "all" TUs, just housekeeping ("hk"), and HUVEC- or GM12878-specific ones. **(i)** HSA 14 in HUVEC. **(ii)** HSA 18 in HUVEC. **(iii)** HSA 19 in HUVEC. **(iv)** HSA 14 in GM12878.

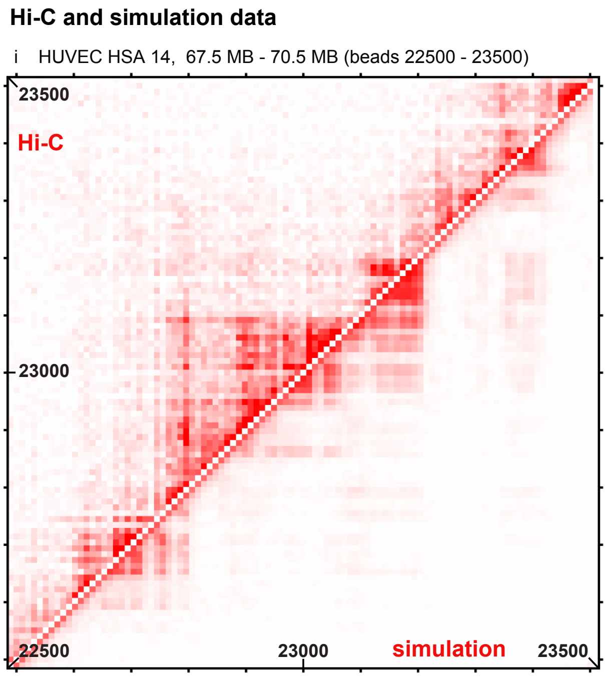


FIG. S5. **Comparing experimental Hi-C and numerical contact maps for HUVEC HSA** 14**.** After running 100 simulations for HUVEC HSA 14, contact maps for two chromosome fragments are generated and compared to an experimental Hi-C map provided by ENCODE [6]. **(i)** fragement $67.5 - 70.5$ MB (beads $22500 - 23500$). **(ii)** fragment $100.8 - 106.8$ MB (beads $33600 - 35500$). In both cases the binning is 30kbp, each pixel is colored according to the number of contacts and fair data agreement is found (Pearson coefficient $r \sim 0.7$, $p$-value $< 10^{-6}$ 2-sided t-test). The diagonal is made white for graphical clarity.

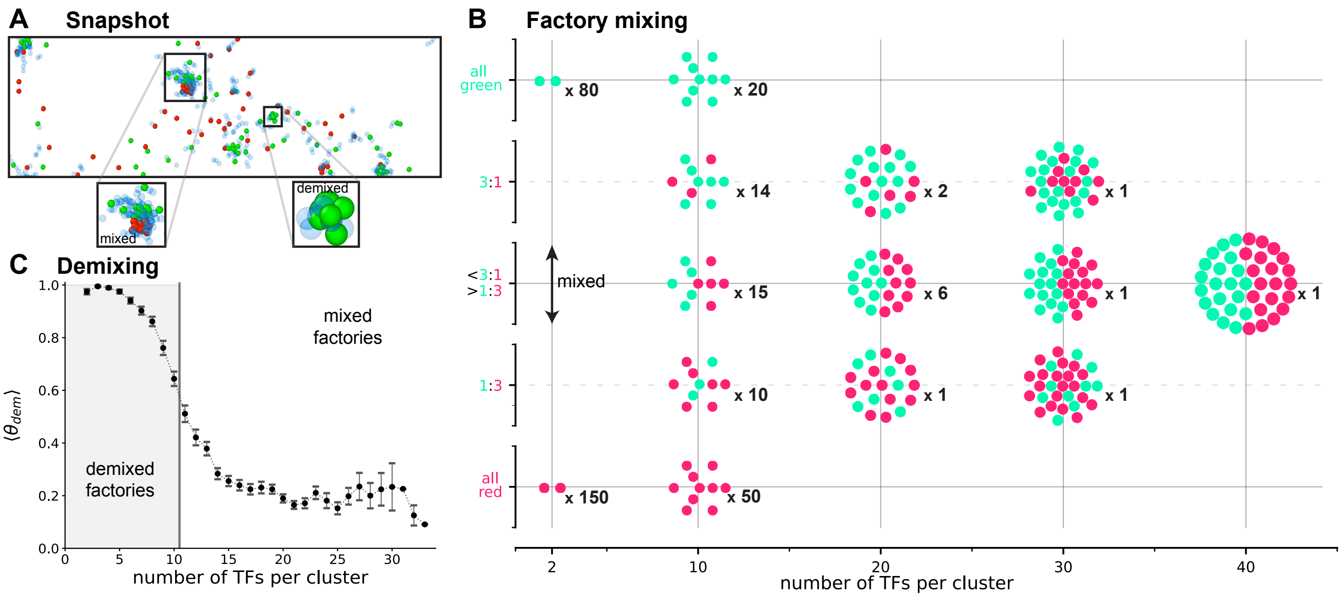


FIG. S6. **Cluster analysis for HSA** 14 **in GM12878.** After running 100 simulations, clusters are identified. **(A)** Snapshot of a typical final conformation (TUs, non-binding beads, and TFs in off state not shown). Insets: a large mixed cluster and a small demixed one. **(B)** Cartoons showing representative contents of clusters (for a more complete representation, see Fig. S7). We consider 5 types of clusters (ranging from all green to all red, $y$ axis), and 5 values for the total numbers of TFs/cluster (2, 10, 20, 30, 40, $x$ axis). Cartoons: cluster composition. Black numbers: observed number of clusters in the simulation. **(C)** Average of the demixing coefficient $\langle\theta_{\rm dem}\rangle$ as a function of cluster size (error bars: SD). Grey area: demixed regime (corresponding to specialized factories) where $\langle\theta_{\rm dem}\rangle$ is $> 0.5$. White area: mixed regime (corresponding to mixed factories associated with HOTs) where $\langle\theta_{\rm dem}\rangle$ is $< 0.5$.

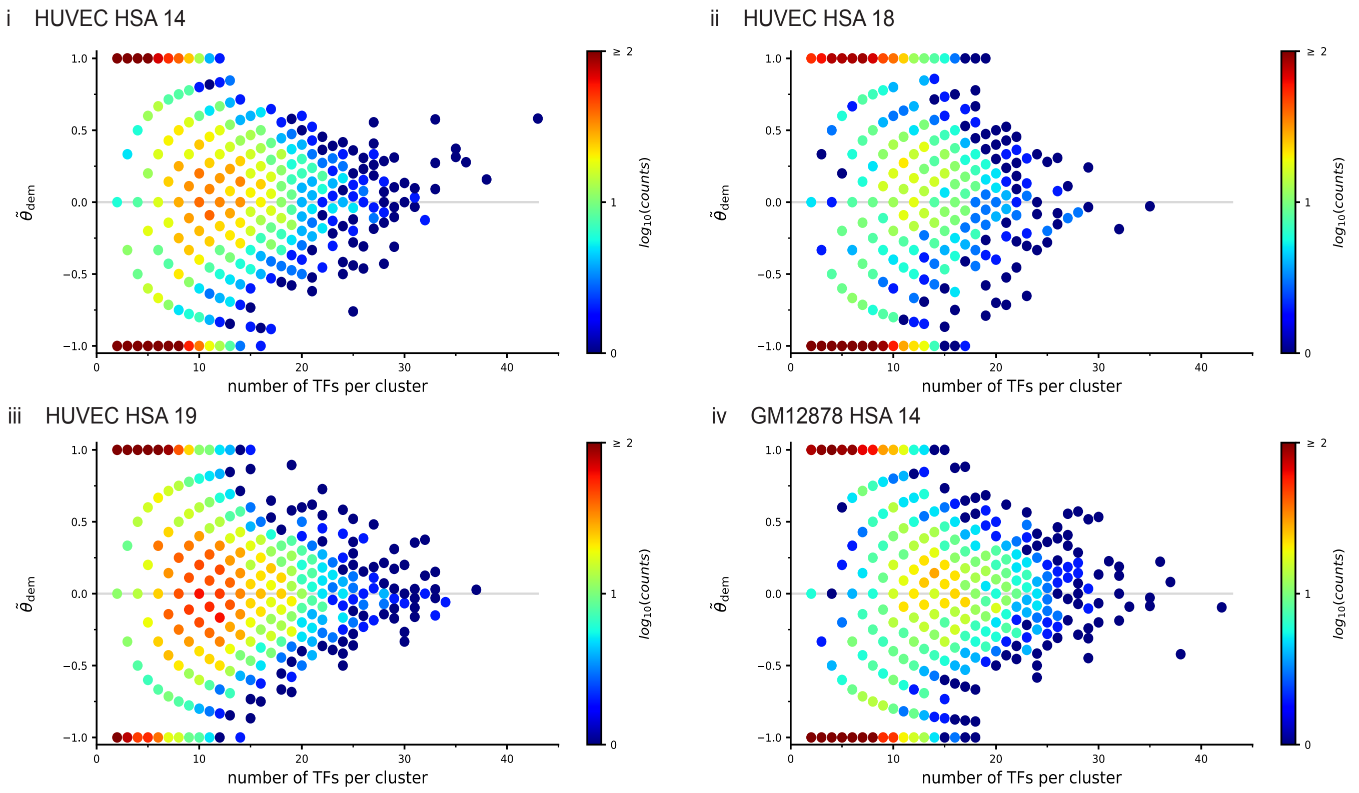


FIG. S7. **Small clusters tend to be less mixed than large ones in simulations of various human chromosomes in two cell types.** After running 100 simulations for each chromosome in each cell type, all clusters are identified, numbers of housekeeping and cell-type-specific TUs in them counted, and demixing coefficients determined. Each dot in a plot is positioned according to TF content and value of $\tilde{\theta}_{\rm dem} \equiv = (2x_{\rm red} - 1)$, with $x_{\rm red}$ the fraction of red beads in a cluster (so that $\tilde{\theta}_{\rm dem} = +1$ $\tilde{\theta}_{\rm dem} = -1$ correspond to fully demixed red and green clusters respectively ). Each dot is coloured according to its frequency of occurrence (right-hand bars). **(i)** HSA 14 in HUVEC. **(ii)** HSA 18 in HUVEC. **(iii)** HSA 19 in HUVEC. **(iv)** HSA 14 in GM12878. All panels have similar patterns; each indicates there are high numbers of demixed small clusters (top and bottom left of each panel), and lower numbers of mixed larger clusters (at middle right in each panel).

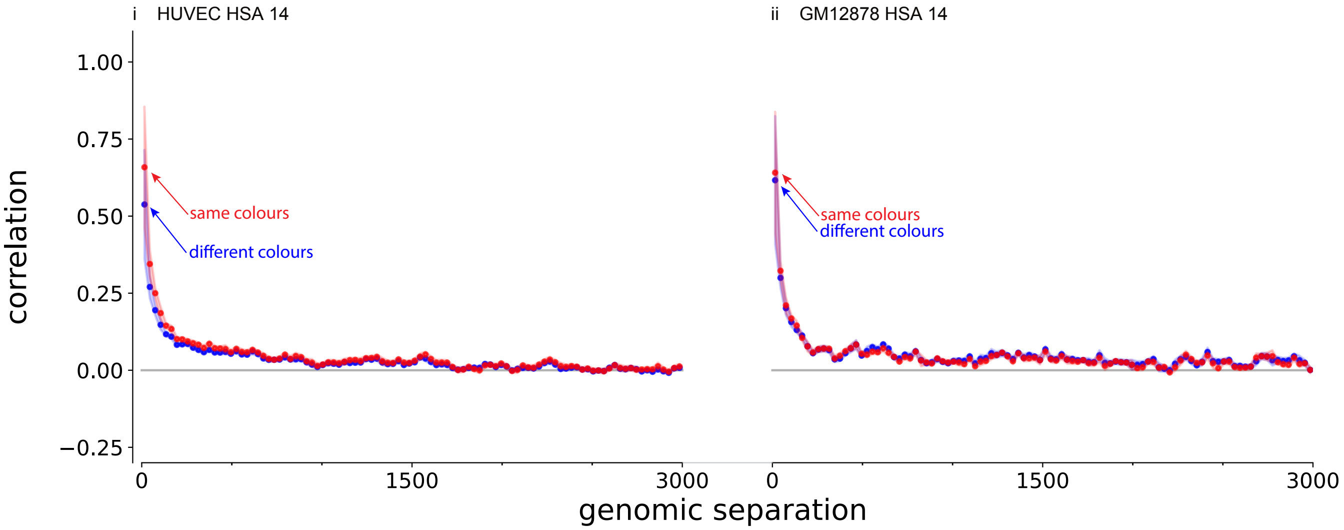


FIG. S8. **Effect of genomic separation on activity correlations found in whole human chromosomes.** Correlation values (shading shows +/-SD, and is usually less than line/spot thickness) at fixed genomic distance are taken from super-/sub-diagonals of Pearson matrices. Red dots give mean correlations between TUs of same colour, and blue dots those between TUs of different colours. **(i)** HSA 14 in HUVEC. **(ii)** HSA 14 in GM 12878.

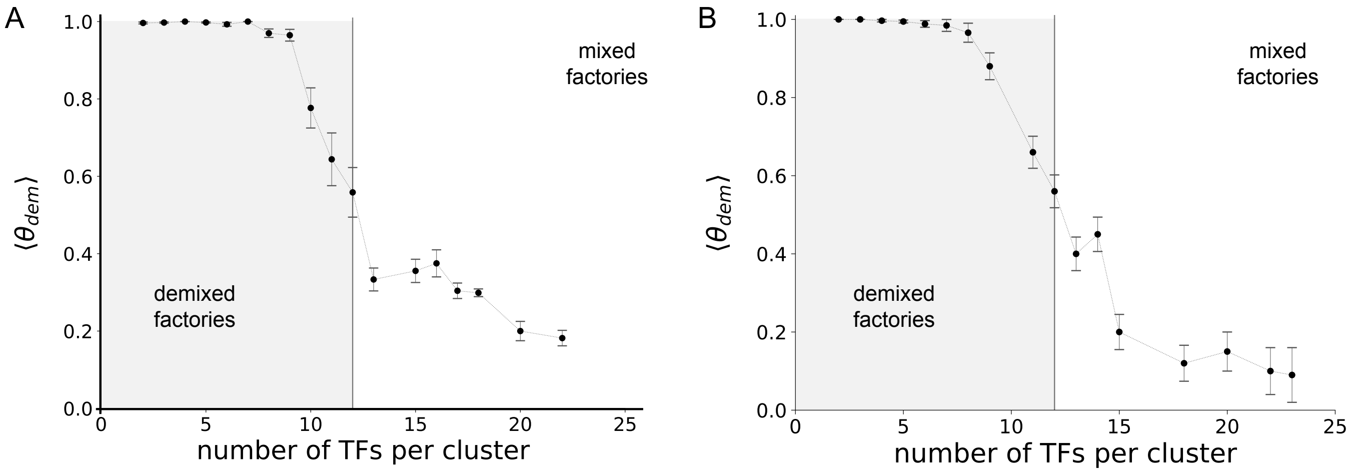


FIG. S9. **Cluster analysis for HSA 14 in HUVEC with smaller TFs.** Clusters are identified for HSA 14 in HUVEC after running 80 simulations with TFs size $0.5\sigma$ **(A)** and 100 simulations with TFs size $0.16\sigma$ **(B)**. Panels report average of the demixing coefficient $\langle\theta_{dem}\rangle$ as a function of cluster size (error bars: SD). Grey area: demixed regime (corresponding to specialized factories) where $\langle\theta_{dem}\rangle > 0.5$. White area: mixed regime (corresponding to mixed factories associated with HOTs) where $\langle\theta_{dem}\rangle < 0.5$.

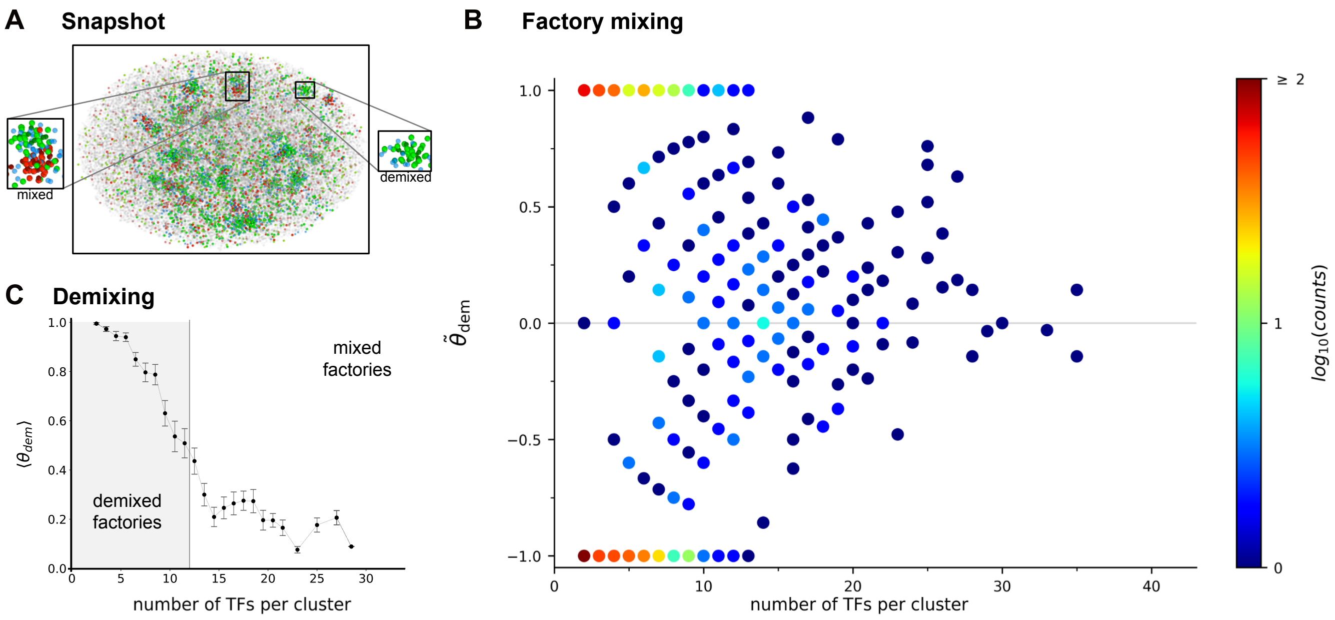


FIG. S10. **Cluster analysis for confined HSA 14 in HUVEC.** After running 100 simulations, clusters are identified for HSA 14 in HUVEC. The system is confined into an ellipsoidal territory with aspect ratio chosen according to typical experimental values [20] (semiaxes were $22.24\sigma : 34.24\sigma : 41.80\sigma$, or $0.67\mu m : 1.03\mu m : 1.25\mu m$. Confinement is enforced by modifying the source code in LAMMPS to describe an ellipsoidal indenter. This introduces a soft force towards the centre of the ellipsoid, only if beads exit the confining ellipsoid. **(A)** Snapshot of a typical confined conformation. Insets: a large mixed cluster and a small demixed one (non-binding beads and TFs in off state not shown). **(B)** All clusters are identified, numbers of housekeeping and cell-type-specific TUs in them counted, and demixing coefficients determined. Each dot in the plot is positioned according to TF content and value of $\tilde{\theta}_{dem}$ and coloured according to its frequency of occurrence (right-hand bar). **(C)** Average of the demixing coefficient $\langle\theta_{dem}\rangle$ as a function of cluster size (error bars: SD). Grey area: demixed regime where $\langle\theta_{dem}\rangle > 0.5$. White area: mixed regime where $\langle\theta_{dem}\rangle < 0.5$.

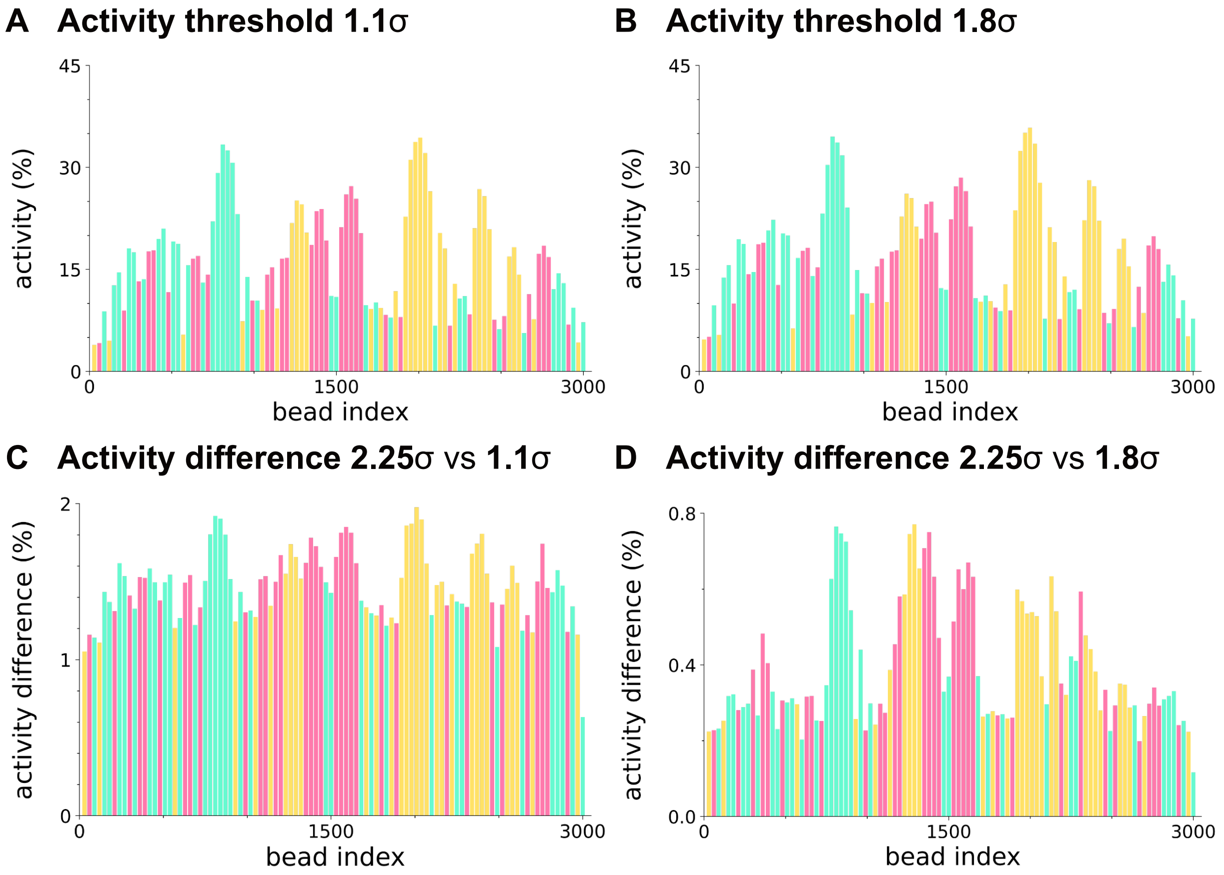


FIG. S11. **Varying transcriptional activity threshold does not alter transcriptional profiles.** **(A)** and **(B)** Bar heights give transcriptional activities of each TU in the toy string (average of 100 runs each lasting 8 $10^5\tau_B$). A TU bead is considered to be active respectively whilst within $1.8\sigma \sim 5.4 \times 10^{-8}m$ and $1.1\sigma \sim 3.3 \times 10^{-8}m$ of a TF:pol complex of similar colour. **(C)** and **(D)** Difference between the transcriptional profile obtained with threshold $2.25\sigma$ (see Fig.1D) and those obtained with threshold $1.8\sigma$ and $1.1\sigma$, respectively.

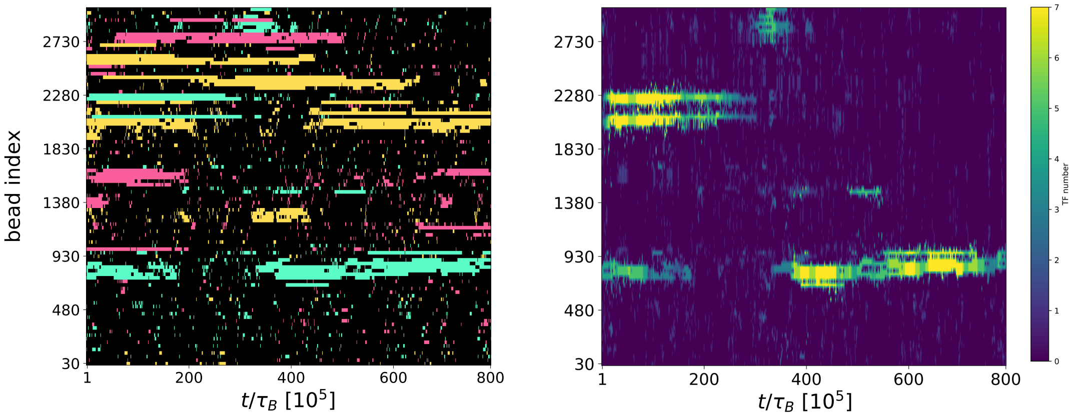


FIG. S12. Left: Kymograph showing, at each time step, the transcriptionally active TUs, color-coded according to their assigned type. This kymograph is identical to that reported in Fig. 2E(i) of the main text. Right: Corresponding kymograph displaying the number of green TFs within a sphere of radius $10\,\sigma$ centered on each green TU.

[1] C. A. Brackey, D. Marenduzzo, and N. Gilbert, “Mechanistic modeling of chromatin folding to understand function,” *Nature Methods*, vol. 17, no. 8, 2020. [Online]. Available: https://doi.org/10.1038/s41592-020-0852-6

[2] C. A. Brackley, S. Taylor, A. Papantonis, P. R. Cook, and D. Marenduzzo, “Nonspecific bridging-induced attraction drives clustering of DNA-binding proteins and genome organization,” *Proceedings of the National Academy of Sciences USA*, vol. 110, no. 38, pp. E3605–E3611, 2013.

[3] A.-M. Florescu, P. Therizols, and A. Rosa, “Large scale chromosome folding is stable against local changes in chromatin structure,” *PLOS Computational Biology*, vol. 12, pp. 1–21, 2016.

[4] C. Brackley, N. Gilbert, D. Michieletto, A. Papantonis, M. Pereira, P. Cook, and D. Marenduzzo, “Complex small-world regulatory networks emerge from the 3d organisation of the human genome,” *Nat. Commun.*, vol. 12, no. 1, pp. 1–14, 2021.

[5] A. P. Thompson, H. M. Aktulga, R. Berger, D. S. Bolintineanu, W. M. Brown, P. S. Crozier, P. J. in ’t Veld, A. Kohlmeyer, S. G. Moore, T. D. Nguyen, R. Shan, M. J. Stevens, J. Tranchida, C. Trott, and S. J. Plimpton, “LAMMPS - a flexible simulation tool for particle-based materials modeling at the atomic, meso, and continuum scales,” *Comp. Phys. Comm.*, vol. 271, p. 108171, 2022.

[6] ENCODE project portal. [Online]. Available: https://www.encodeproject.org

[7] A. R. Quinlan and N. Kindlon. bedtools: a powerful toolset for genome arithmetic. [Online]. Available: https://bedtools.readthedocs.io/en/latest/

[8] E. B. Stovner. epic: find diffusely enriched domains in chip-seq data. [Online]. Available: https://bioepic.readthedocs.io/en/latest/

[9] Y. Zhang, T. Liu, C. A. Meyer, J. Eeckhoute, D. S. Johnson, B. E. Bernstein, C. Nusbaum, R. M. Myers, M. Brown, W. Li, and X. S. Liu, “Model-based analysis of chip-seq (MACS),” *Genome Biology*, vol. 9, p. R137, 2008.

[10] E. project. bedgraphtobigwig file format conversion. [Online]. Available: https://github.com/ENCODE-DCC/kentUtils

[11] hg19.chrom.sizes file. [Online]. Available: http://hgdownload.cse.ucsc.edu/goldenpath/hg19/bigZips/hg19.chrom.sizes

[12] A. Buckle, C. A. Brackley, S. Boyle, D. Marenduzzo, and N. Gilbert, “Polymer simulations of heteromorphic chromatin predict the 3-d folding of complex genomic loci,” *Molecular Cell*, vol. 72, pp. 786–797, 2018.

[13] G. Fudenberg, M. Imakaev, C. Lu, A. Goloborodko, N. Abdennur, and L. Mirny, “Formation of chromosomal domains by loop extrusion,” *Cell Reports*, vol. 15, no. 9, pp. 2038–2049, 2016.

[14] A. Goloborodko, J. Marko, and L. Mirny, “Chromosome compaction via active loop extrusion,” *Biophysical Journal*, vol. 110, pp. 2162–2168, 2016.

[15] E. J. Banigan and L. A. Mirny, “Loop extrusion: theory meets single-molecule experiments,” *Current Opinion in Cell Biology*, vol. 64, pp. 124–138, 2020, cell Nucleus.

[16] P. R. Cook and D. Marenduzzo, “Transcription-driven genome organization: a model for chromosome structure and the regulation of gene expression tested through simulations,” *Nucleic acids research*, vol. 46, no. 19, pp. 9895–9906, 2018.

[17] S. Boyle, I. M. Flyamer, I. Williamson, D. Sengupta, W. A. Bickmore, and R. S. Illingworth, “A central role for canonical prc1 in shaping the 3d nuclear landscape,” *Genes & development*, vol. 34, no. 13-14, pp. 931–949, 2020.

[18] A. R. Strom, A. V. Emelyanov, M. Mir, D. V. Fyodorov, X. Darzacq, and G. H. Karpen, “Phase separation drives heterochromatin domain formation,” *Nature*, vol. 547, no. 7662, pp. 241–245, 2017.

[19] C. A. Brackley, B. Liebchen, D. Michieletto, F. Mouvet, P. R. Cook, and D. Marendu zzo, “Ephemeral protein binding to DNA shapes stable nuclear bodies and chromatin domains,” *Biophysical Journal*, vol. 112, no. 6, pp. 1085–1093, 2017.

[20] Y. Wang, M. Nagarajan, C. Uhler, and G. V. Shivashankar, “Orientation and repositioning of chromosomes correlate with cell geometry-dependent gene expression.” *Mol Biol Cell*, vol. 28, no. 14, pp. 1997–2009, Jul 2017.
